# Resolving inconsistent effects of tDCS on learning using a homeostatic structural plasticity model

**DOI:** 10.1101/2025.01.23.634520

**Authors:** Han Lu, Lukas Frase, Claus Normann, Stefan Rotter

## Abstract

Transcranial direct current stimulation (tDCS) is increasingly used to modulate motor learning. Current polarity and intensity, electrode montage, and application before or during learning had mixed effects. Both Hebbian and homeostatic plasticity were proposed to account for the observed effects, but the explanatory power of these models is limited. In a previous modeling study, we showed that homeostatic structural plasticity (HSP) can explain long-lasting after-effects of tDCS and transcranial magnetic stimulation (TMS). The interference between motor learning and tDCS, which are both based on HSP in our model, is a candidate mechanism to resolve complex and seemingly contradictory experimental observations. We implemented motor learning and tDCS in a spiking neural network subject to HSP. The anatomical connectivity of the engram induced by motor learning was used to quantify the impact of tDCS on motor learning. Our modeling results demonstrated that transcranial direct current stimulation applied before learning had weak modulatory effects. It led to a small reduction in connectivity if it was applied uniformly. When applied during learning, targeted anodal stimulation significantly strengthened the engram, while targeted cathodal or uniform stimulation weakened it. Applied after learning, targeted cathodal, but not anodal, tDCS boosted engram connectivity. Strong tDCS would distort the engram structure if not applied in a targeted manner. Our model explained both Hebbian and homeostatic phenomena observed in human tDCS experiments by assuming memory strength positively correlates with engram connectivity. This includes applications with different polarity, intensity, electrode montage, and timing relative to motor learning. The HSP model provides a promising framework for unraveling the dynamic interaction between learning and transcranial DC stimulation.

## Introduction

Transcranial direct current stimulation (tDCS) is a promising non-invasive brain stimulation with a long history [1, 2, 3]. By applying a weak direct current (1 – 2 mA) to the brain through external surface electrodes attached to the scalp, although not demonstrated in all studies, tDCS can instantly modulate the firing activity of active neurons by influencing the release of vesicles from presynaptic sites [4, 5, 6, 7] or polarizing the membrane potential [8, 9, 10, 11, 12, 13, 14] and influencing network activity [15]. However, changes in neural excitability and behavioral effects persist up to hours or even days after stimulation has been terminated [16, 17, 18, 19]. Exploiting these direct and indirect effects, tDCS is increasingly used in experimental and clinical settings to modulate brain functions [20, 21, 22, 23, 24, 25] and treat diseases [14, 26, 27, 28, 29, 30, 31, 32, 33]. With increased usage and growing diversity of tDCS studies, it is now often applied in patients and healthy volunteers that are concurrently engaged in other tasks, such as learning [25, 28, 34, 35, 36, 37, 38, 39].

Motor learning is the paradigm most frequently studied in tDCS, as there are established protocols and effective tests for cortical excitability [28, 40, 41, 42], as well as its great potential in rehabilitating motor skills with low adverse events in patients with Parkinson’s disease or after stroke [43, 44, 45, 46, 47, 48, 49], etc. Among these studies, the cerebellum, the primary motor cortex (M1), the premotor cortex (M2), the prefrontal cortex (PFC), and the posterior parietal cortex (PPC) are the most targeted area, where parameters such as current polarity, current intensity, electrode montage, and stimulation timing are acknowledged to have a discernible influence on the outcome of stimulation (see a recent review in [50]). For example, when applying tDCS to M1 concurrently with a motor learning task, it was reported that anodal tDCS improved learning [51, 52], while cathodal tDCS did not produce comparable effects [42, 53] or even reduced retention [54]. However, some follow-up studies that used slightly different configurations or tasks failed to replicate these effects [28, 41, 55, 56]. In addition, the strength of the after-effects on cortical excitability did not correlate well with the current intensity [57, 58, 59].

The focality of stimulation depends on the electrode montage, and this is known to also influence the effect of tDCS on motor learning. Traditionally, either unilateral (with the anode on the non-dominant M1 and the cathode over the contralateral supraorbital area) or bilateral (anode over the dominant and cathode over the non-dominant M1) electrode montages are used. Interestingly, these two configurations produce different results [60, 61]. To improve focality in a specific brain area, high definition tDCS (HD-tDCS) was proposed, *e.g.* employing a central anode surrounded by four cathodes. HD-tDCS induces a more focal electric field that was shown in recent pilot studies to facilitate motor learning [62, 63] and differentially recruits different pathways to M1 [63]. However, as more and more studies emerge, it becomes clear that the relation between motor learning and tDCS strongly depends on the learning phase. The same tDCS protocol applied before learning, during training blocks, or after learning leads to very different results [39, 64, 65, 66, 67]. These effects appear to differ in brain area [39], subject age [38], and health condition [68]. Assuming that tDCS is, in principle, a perturbation of the underlying neural activity, the outcome of tDCS should be brain-state dependent. However, the neural correlate of this dependency has not yet been systematically elucidated.

Synaptic plasticity was recognized as the key mechanism underlying the modulatory effects of tDCS on motor learning. Animal studies focused primarily on M1 and suggested that M1 displays strong activity-dependent plasticity [69], and dendrite-specific spine formation was observed in motor skills training [70, 71]. After three days of *in vivo* tDCS stimulation (20 min per day) in M1, mice presented improved synaptic transmission and increased spine density in pyramidal neurons in layer II / III of M1, as well as improved motor performance [72]. Indirect evidence from human experiments applying tDCS in motor learning is abundant, where both Hebbian and homeostatic phenomena have been reported [73]: For example, pre-learning anodal tDCS was considered to trigger homeostatic inhibition and hinder learning performance. In contrast, tDCS concurrent with learning leads to boosting effects similar to long-term potentiation (LTP). This was interpreted as a Hebbian phenomenon. However, the explanatory power of currently used models of homeostatic and Hebbian plasticity is limited [74]. In addition, those models do not account for spine turnover. We recently showed that the homeostatic structural plasticity model (HSP) can provide a theoretical framework to explain the long-lasting after-effects of tDCS [75], repetitive transcranial magnetic stimulation (rTMS) [76], and repetitive optogenetic stimulation [77]. According to the HSP rule, neurons continuously grow synaptic elements (boutons and spines) and specifically form new synapses if their neural activity falls below homeostatic *set-point* [75, 77, 78, 79, 80]. The HSP rule can also implicitly have associative properties (Hebbian-like) on top of its primary function to homeostatically regulate neural activity [79, 80]. In line with this idea, the HSP model is considered a good candidate to explain the dynamic interaction between tDCS and motor learning.

In our current study, we conceived motor learning as the formation of new structures corresponding to new memories (“engrams”) in a recurrent network. Transcranial stimulation was assumed to alter the equilibrium between neuronal activity and network structure, leading to additional network rewiring that interferes with learning. We analyzed the effect of different combinations of stimulation parameters, such as electrode montage, DC polarity and intensity, as well as the relation to the learning phase. In summary, our model can reconcile the observed impact of DC stimulation on motor learning and reconcile seemingly contradicting experimental results. We also made predictions that have not yet been reported by human experiments. The HSP model, linking structural changes and the homeostatic regulation of neural activity, provides a systematic framework for explaining and predicting the effects of tDCS on motor learning.

## Methods

### Neuron, synapse, and network models

Numerical simulations of neural networks with homeostatic structural plasticity (HSP) were used to explore the interaction between tDCS and learning. We used the same neuron model, synapse model, and network model, as published in our previous paper [75, 80, 81]. The NEST simulator [82] with parallel MPI-based computation was used to perform the large-scale neural network simulations presented here.

We used a leaky integrate-and-fire (LIF) neuron model. The cortical area M1 was conceived as an inhibition-dominated recurrent neural network of 10 000 excitatory and 2 500 inhibitory neurons [83]. All connections involving inhibitory neurons were static with a fixed weight. Recurrent synapses among excitatory neurons (E-E connections) were grown according to the HSP rule [75, 79, 80]. Each neuron in the network received the same Poissonian external input. After the 750 s growth period, the network had established an equilibrium between neural activity and network architecture. Excitatory neurons fired asynchronously and irregularly at 8 Hz and were connected with a connection probability around 9 %. Details concerning neuron and network models, as well as the HSP rule, are described in the Supplementary Materials.

### Motor learning protocol

In a previous work, learning and memory were simulated as an engram [80]. Here we follow the idea and simulate the motor learning training process in a motor engram comprising 1 000 excitatory neurons (10 % of all excitatory neurons) consistently for all experiments. The neurons were driven by Poissonian spike input at a rate of 1.5 kHz with synaptic weight *J* = 0.1 mV for 150 seconds. The input of motor learning leads to a fluctuating membrane potential with a mean value *µ* = *ν_ext_τ_m_J*_learning_ = 1.50 mV [83]. The connectivity of the motor engram (Γ_engram_) was conceived as the main readout that represents the strength of memory.

### Direct current stimulation protocol

A model of how tDCS affects single neurons was published and characterized in our previous paper [75]. Thus, we assume that the electric field induced by the administration of transcranial current polarizes the soma of neurons with extended nonisotropic morphology, such as pyramidal neurons [13, 14]. The duration of the stimulus was 150 s for all experiments. However, we tested different configurations for the polarity and intensity of the current and of the electrode montage.

In all of our experiments, the E-E connections were grown during an initial 750 s growth period in the presence of background input. The motor learning protocol and/or tDCS were applied after the network had reached its structural equilibrium.

#### Electrode montage

We first devised three paradigmatic scenarios to explore the impact of the electrode montage. The traditional two-electrode montage may induce a diffuse electric field that covers a volume larger than the volume engaged in motor learning. In contrast, the HD-tDCS montage induces a focal electric field that targets only a subset of neurons. In our current study, we concentrated on three extreme scenarios: uniform, targeted, and unfocused. In the uniform scenario, DC stimulation was applied homogeneously to all excitatory neurons in the network. Targeted DC stimulation was administered exclusively to the motor engram. Unfocused stimulation was an intermediate scenario, in which DC stimulation covers only half of the engram cells and the same amount of non-engram excitatory neurons.

#### Relative timing of motor learning and transcranial DC stimulation

We tested three experimental conditions for each electrode montage. We applied tDCS immediately before, during, or immediately after learning input.

#### Polarity and intensity of transcranial DC

For each combination of electrode montage and relative timing, we systematically varied the amplitude of tDCS to polarize the membrane potential between *−*0.6 mV and 0.6 mV. If not stated otherwise, the transcranial DC currents used in this study were generally weaker than the motor learning input (1.5 mV).

### Measurement of firing rate and connectivity

To quantify the modulatory effect of tDCS on learning, we considered the temporal evolution of neural activity and the connectivity of the motor engram. The neural firing rate was calculated from the spike count in a recording window of 5 s. The mean activity of the population was estimated by averaging the firing rates of individual neurons. We used a *n × n* connectivity matrix (*A_ij_*) to represent the recurrent excitatory connections of our network, where columns and rows correspond to pre- and postsynaptic neurons. The entry *A_ij_* of the matrix represents the total number of synaptic connections from neuron *j* to neuron *i*. The mean network connectivity at any given time *t* was calculated by 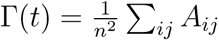.

## Results

### Modeling motor learning with a motor engram

We modeled motor learning by introducing excitatory spike input to a subgroup of the excitatory neurons (Figure 1A-B, purple shading). The application and termination of motor learning input substantially altered the neural activity of stimulated neurons and induced homeostatic reorganization of synapses as previously described [75, 79, 80]. When the disrupted neural activity returned to the homeostatic level, the network architecture did not recover to the pre-learning state. Instead, neurons receiving motor learning input wired more with each other, but less with other non-stimulated excitatory neurons (Figure 1C). This cell assembly with elevated connectivity is referred to as a motor engram in the following.

**Figure 1:**
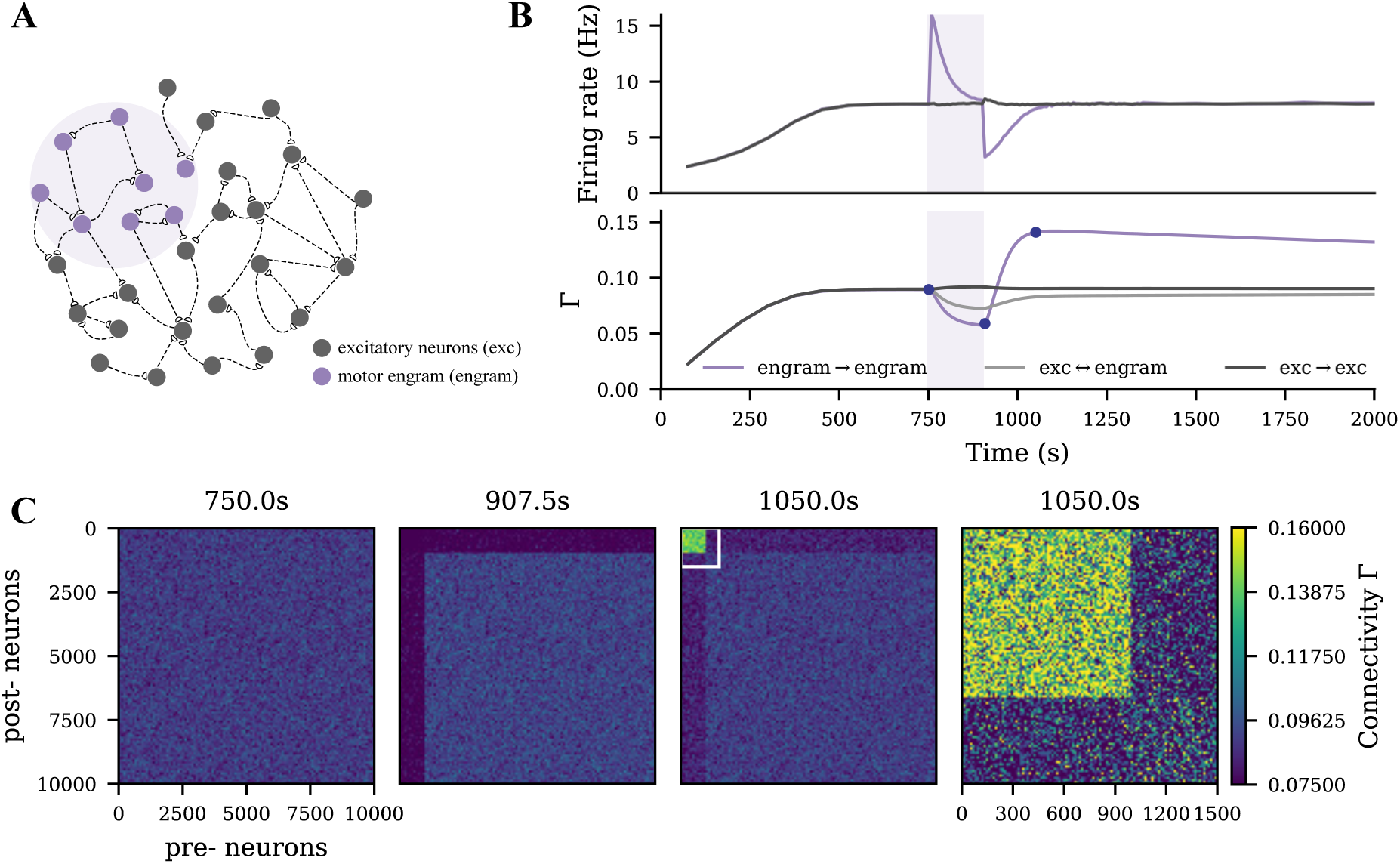
Strong motor engram formed during motor learning. **A** Schematic of the M1 neural network, part of which (10 %) was subject to input that drive motor learning. **B** Temporal evolution of neural activity and network connectivity (Γ) during a typical motor learning episode (purple shading). The purple and dark gray curves in the upper panel represent the firing rates of the motor engram and of the remaining excitatory neurons, respectively. The purple, dark gray, and light gray curves in the lower panel represent the intra-group connectivity of the memory engram, intra-group connectivity of the nonstimulated excitatory neurons, and inter-group connectivity between both populations. **C** The E-E connectivity matrix at the three time points indicated in panel **B** (dark blue dots), the fourth panel shows a scaling of motor-engram connectivity. The color scale accounts for connectivity (connection probability).

### tDCS triggers similar but weaker network remodeling than motor learning

We modeled tDCS by injecting subthreshold direct current into the soma as described in our previous study [75]. When the membrane potential of a subpopulation was depolarized or hyperpolarized by DC (Figure 2A-C, green shading), a similar synapse reorganization process occurred. The DC input strength that we used in the current study was relatively weaker than the learning input, so the connectivity of the tDCS-stimulated cell assembly was accordingly weaker than the motor engram (Figure 2D-E). In contrast, no resulting cell assembly was triggered when tDCS was applied globally to all excitatory neurons.

**Figure 2:**
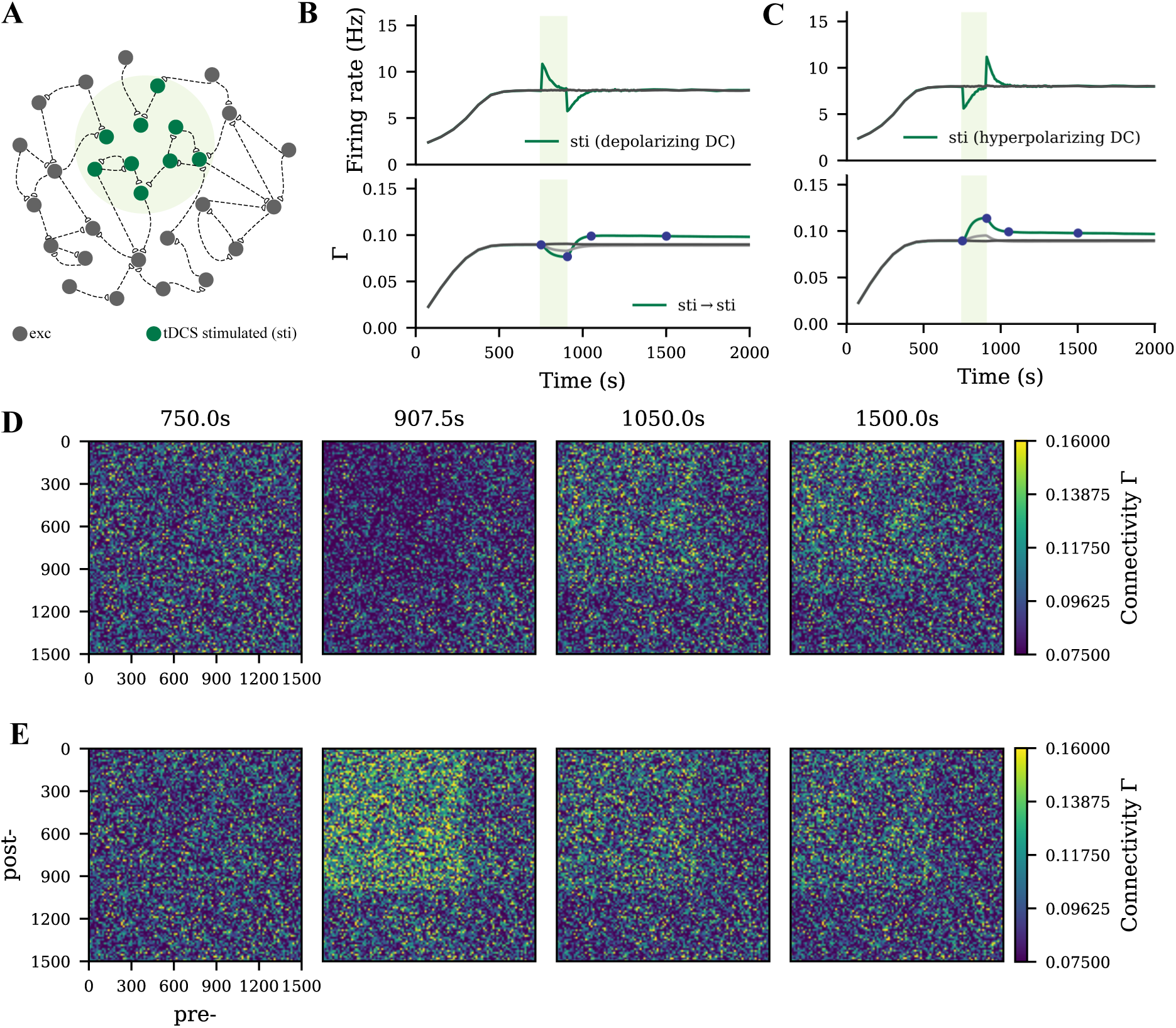
Weak cell assembly formed during tDCS. **A** Schematic of the M1 neural network, part of which (10 %) was weakly polarized by the electrical field applied transcranially. **B-C** Temporal evolution of neural activity and network connectivity, when the membrane potential of neurons is depolarized or hyperpolarized during tDCS. **D-E** The E-E connectivity matrix at four time points indicated in panels **B** and **C** (dark blue dots). Depolarizing and hyperpolarizing tDCS triggered different processes of synaptic reorganization, but both lead to a similar persistent increase in connectivity among stimulated neurons.

### Uniform DC stimulation weakens the motor engram

To capture a rather diffusive tDCS scenario, uniform tDCS was administered to all excitatory neurons before, during or after the learning process, with varying polarity and intensity. In Figure 3B-C, we displayed the evolution of the firing rate and connectivity of the motor engram for all conditions. When applied before the learning process, uniform anodal tDCS slightly increased motor engram connectivity, while cathodal stimulation led to a reduction in connectivity. However, when applied concurrently with learning or post-learning, the uniform tDCS weakened the motor engram irrespective of the DC polarity. The weakening effects further depended on the DC intensity: Stronger tDCS resulted in lower connectivity of the motor engram. In summary, our simulation results demonstrated a divergent impact of uniform tDCS on learning, but with a strong tendency to negatively influence motor learning.

**Figure 3:**
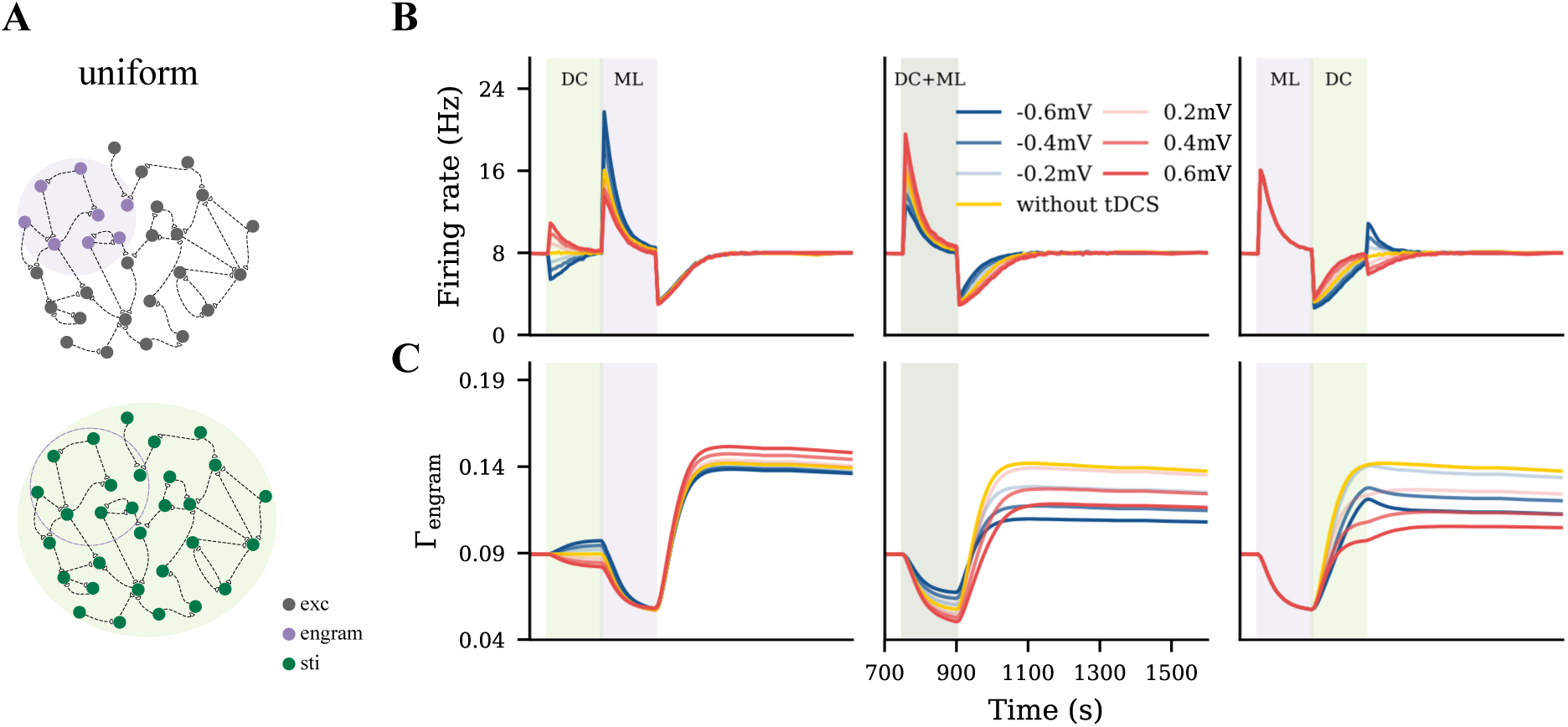
Uniform tDCS generally hindered motor learning. **A** Uniform tDCS was applied to the entire M1 network where the motor engram is embedded. **B-C** Temporal evolution of firing rate and connectivity under three different conditions: applying uniform tDCS immediately before, during, or immediately after learning input. The red and blue curves represent depolarizing and hyperpolarizing DCs, respectively. The yellow curve represents the condition without tDCS. The green and purple shaded areas represent the period of tDCS application (DC) and motor learning (ML), respectively.

### Targeted tDCS influences motor engram in a phase- and polarity-dependent manner

To model a targeted focal stimulation, we applied tDCS exclusively to excitatory neurons within the motor engram before, during, and after motor learning with different polarity and intensity. Our simulation results showed that the final connectivity of the motor engram strongly depended on the relative phase of motor learning and the polarity of tDCS (Figure 4). When applied before motor learning, anodal and cathodal-centered tDCS slightly facilitated engram connectivity. However, when applied concurrently, anodal DC boosted the engram connectivity, whereas cathodal stimulation weakened it. Furthermore, when applied post-learning, the opposite effects were observed: cathodal DC increased the connectivity of the motor engram while the anodal DC reduced it. In summary, our simulation results suggested that the relative learning phase and polarity of tDCS should be chosen with care; targeted anodal and cathodal tDCS might exert opposite effects at different stages of motor learning.

**Figure 4:**
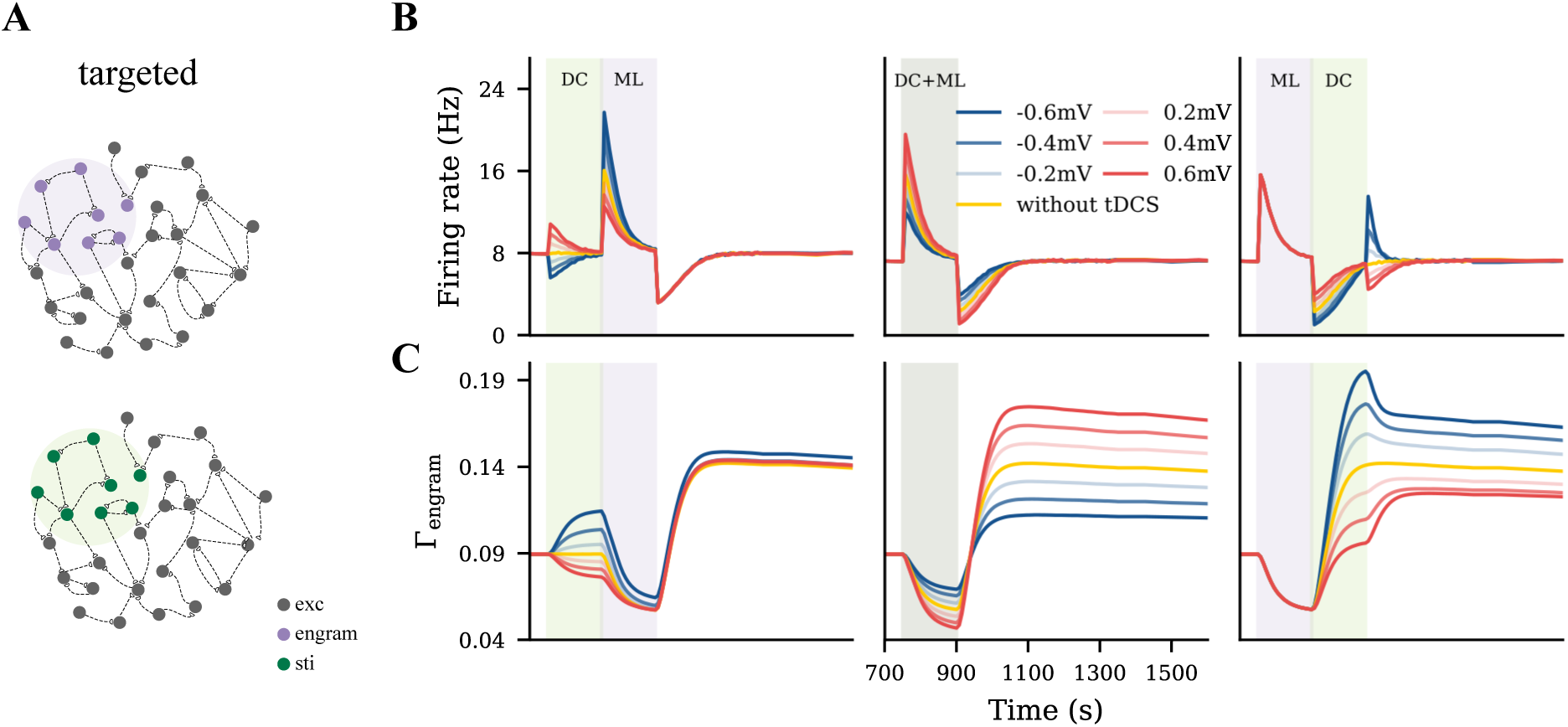
Targeted tDCS exerted opposite effects when applied at different stages of learning. **A** Targeted tDCS matched with the entire motor engram. **B-C** Temporal evolution of the motor engram’s firing rate and connectivity, when targeted tDCS was applied either immediately before, during, or immediately after the learning input.

### Unfocused tDCS and the interaction of cell assemblies

The uniform and targeted tDCS represent two extreme conditions, in which tDCS either homogeneously covers the entire network or exclusively targets the motor engram. Neither holds in the practice of the application of tDCS in humans. So we proposed an intermediate condition, unfocused tDCS, by shifting the targeted tDCS to cover only half of the motor engram (Figure 5A). We expected the motor engram and the stimulated cell assembly to interact with each other during the dynamic stimulation process mediated by homeostatic structural plasticity. To analyze this effect, we presented two examples by applying a depolarizing tDCS at the same intensity before or after the motor learning process. In Figure 5B and 5D, we show the evolution of the firing rate and connectivity of four subpopulations: The overlapped half of the motor engram (orange), the non-overlapped half of the motor engram (purple), neurons receiving tDCS only (green), and the nonstimulated excitatory neurons (dark gray).

**Figure 5:**
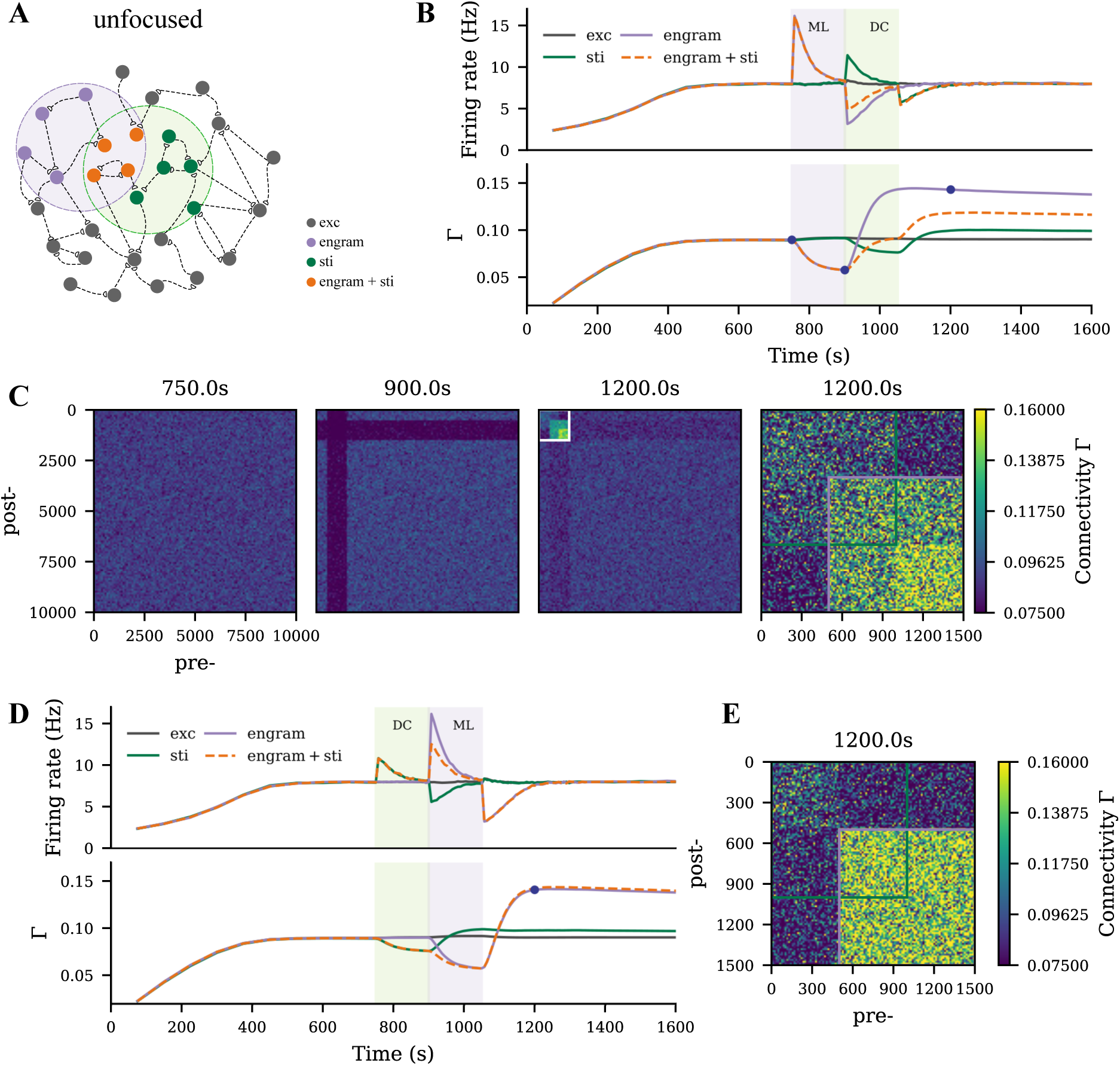
Motor engram and tDCS stimulated cell assembly interfere in case of overlap. **A** Schematic of the M1 neural network, where the motor learning engram and the tDCS stimulated cell assembly had 50 % overlap. **B** Temporal evolution of firing rate and connectivity of excitatory neurons when applying tDCS after learning. The green and purple curves represent neurons that receive only tDCS or only motor learning input, respectively. The orange dashed curve reflects the overlapped half of the engram, and the dark gray curve represents the non-stimulated excitatory neurons. **C** Connectivity matrix at the three time points indicated in panel **B**. In the right panel of **C**, the green square indicates connectivity among the assembly of tDCS-stimulated cells, while the purple square labels the motor learning engram. **D-E** Temporal evolution of neural activity and network connectivity, and the resulting connectivity matrix, when tDCS was applied before motor learning.

When applying the motor learning input prior to tDCS, substantial network remodeling took place to form a motor engram, while the following weak depolarizing tDCS posed the opposite dynamics to the overlapped motor engram. As a result, the connectivity of the overlapped half engram was lower than that of the non-overlapped half (Figure 5C, last panel). In contrast, when applying tDCS before learning, connectivity within the motor engram was not strongly affected, but the overlapped half ended up with higher connectivity than the non-overlapped half (Figure 5E).

To further explore the polarity, intensity, and phase-dependent effects of tDCS in the overlapping scenario, we also performed a systematic analysis. We plotted the temporal evolution of the firing rate and connectivity of the non-overlapped part of the motor engram, the overlapped part of the motor engram, and the tDCS-stimulated cell assembly separately in Figure 6. The non-overlapped half of the engram was slightly affected by tDCS (Figure 6A). The non-overlapped part of the tDCS-stimulated cell assembly was not much influenced by the learning process (Figure 6C). In contrast, the overlapped engram was modulated by tDCS in a contrary way compared to the non-overlapped half (Figure 6B-C). In general, our simulation results showed that weak and strong cell assemblies interacted with each other. The connectivity of the strong cell assembly (motor engram in this case) could be reduced by the following overlapping weak input, due to partial activation, but the connectivity of a weak cell assembly (the tDCS-stimulated cell assembly) is not drastically changed by the following strong input.

**Figure 6:**
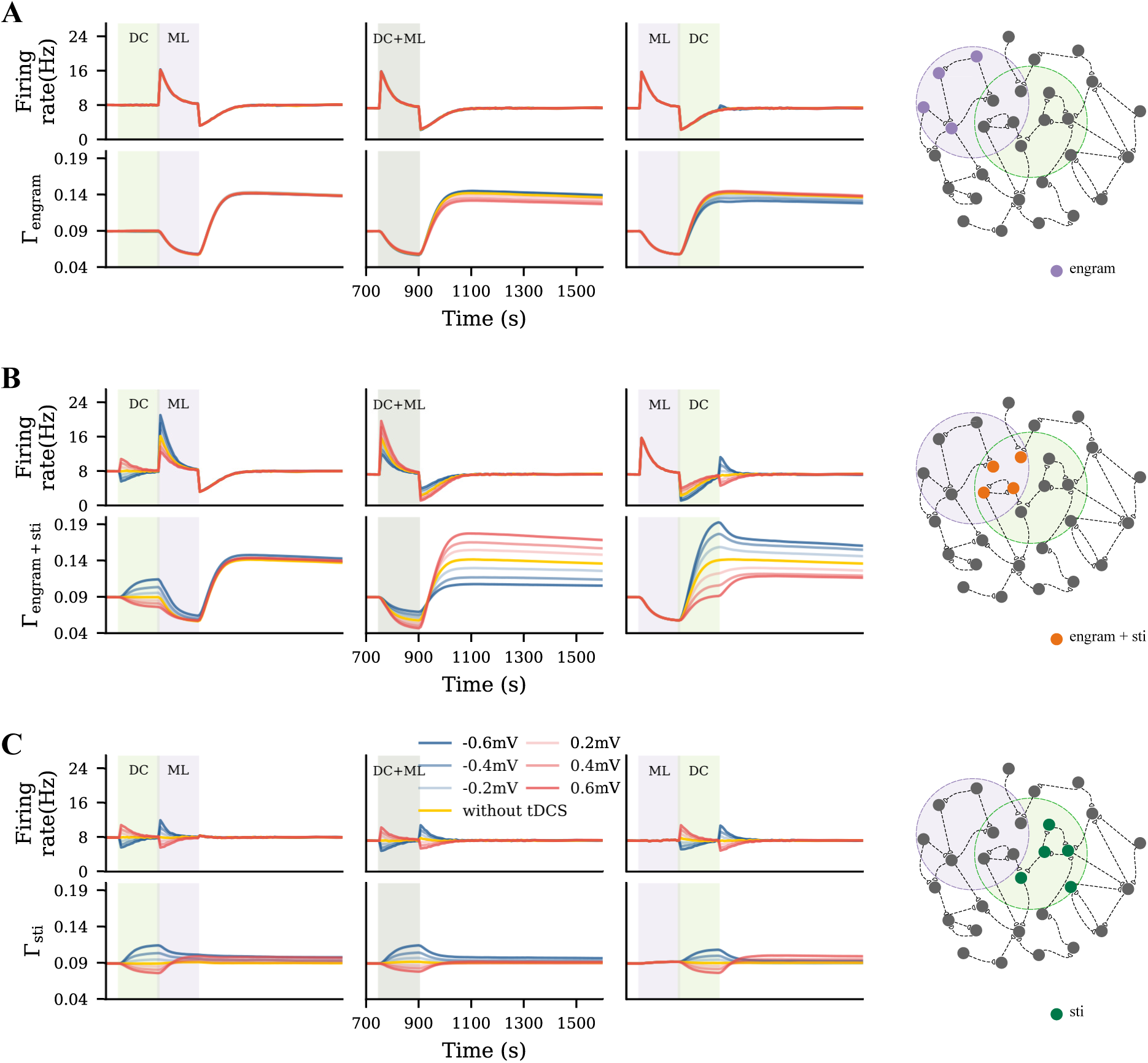
Unfocused tDCS of different intensities applied before, during, or after learning triggered different results. **A-C** Temporal evolution of firing rate and connectivity of the population that only sees motor learning input, the overlapped part of the motor engram that was also stimulated, and the neurons only subjected to tDCS, respectively.

### Strong unfocused tDCS distorted the motor engram

As mentioned above, weak unfocused tDCS modulates motor learning in a phase-dependent manner. Then we wondered if a relatively stronger tDCS, compared to motor learning input, would yield similar results. We used tDCS to achieve a *±*2.8 mV membrane potential and applied it in an unfocused way before or after the motor learning process. As shown in Figure 7A-B, the motor engram at the time when the network reached its equilibrium state (1, 200 s) showed distinct patterns. Compared to the homogeneous motor engram in which no tDCS was applied, strong hyperpolarizing and depolarizing DC—applied either before or after learning—induced local reorganization within the motor engram. The distorted motor engram may not necessarily represent the same learned input regardless of the averaged connectivity.

**Figure 7:**
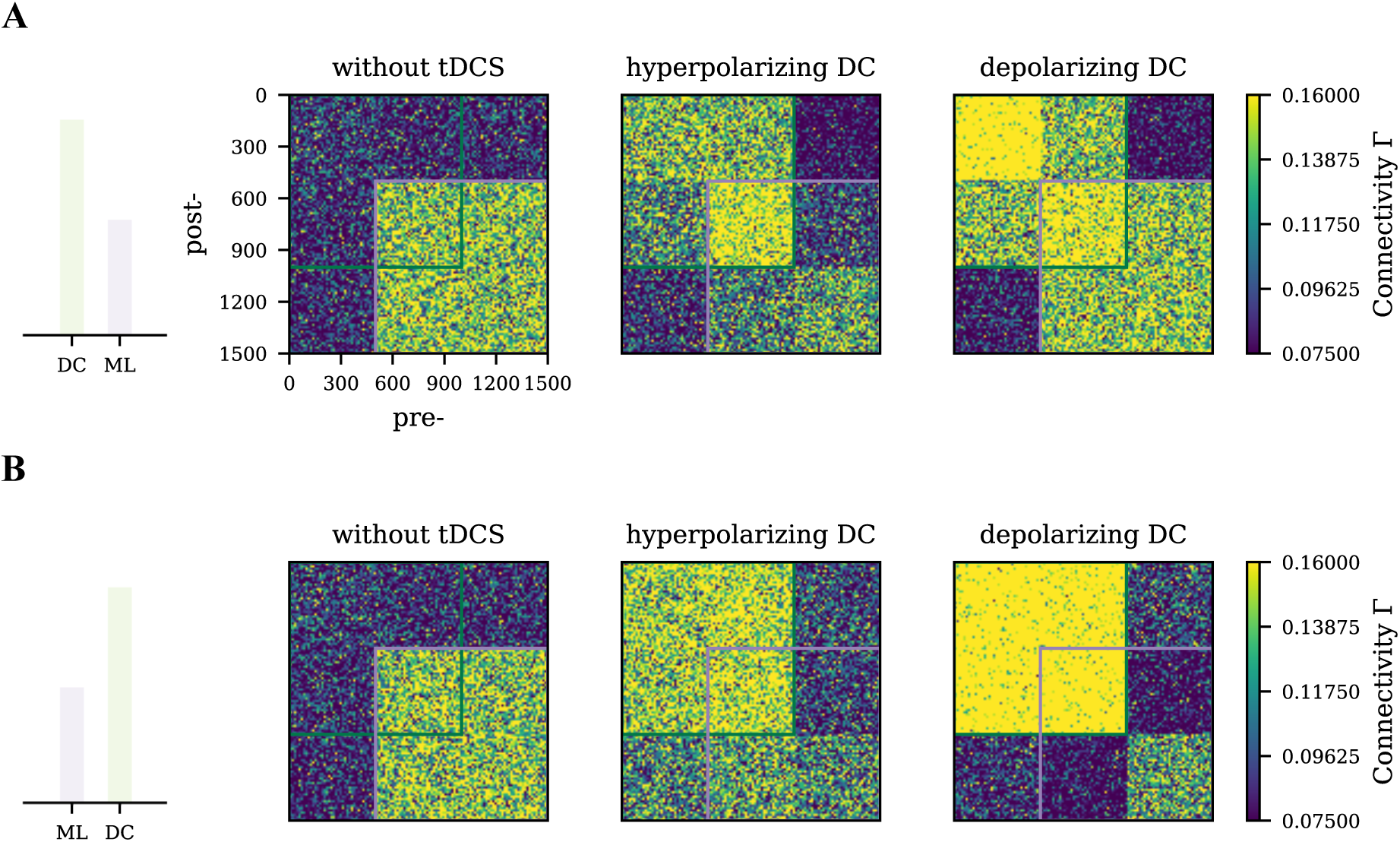
Differential effects of strong tDCS on the motor engram connectivity pattern. Network connectivity was captured at 1, 200 s when the network dynamics reached its equilibrium state. **A** Strong tDCS (hyperpolarizing or depolarizing) was applied before motor learning. The green square labels the connectivity of the tDCS-stimulated cell assembly, while the purple square labels the connectivity of the motor engram. Strong hyperpolarizing and depolarizing tDCS resulted in a different connectivity pattern as compared to the control case where only motor learning input was applied but no tDCS (left panel). **B** Strong tDCS of both polarities was applied after motor learning. Drastic changes in connectivity pattern were observed in the depolarizing tDCS, where the overlapped engram formed a strong connectivity but almost disconnected with the non-overlapped engram.

## Discussion

Motivated by the empirical observations that tDCS triggered changes in both synaptic transmission and dendritic spine density [33, 72], we used a neural network model that implements homeostatic structural plasticity in the present study to systematically explore the interference between tDCS and motor learning. Learning input to the network triggers specific remodeling and induces a memory engram that reflects an altered or entirely new motor program as modeled before [80]. Transcranial DC stimulation also alters the dynamics of the network and induces structural plasticity. However, when applied with different electrode montages and at different phases of motor learning, tDCS interferes with motor learning and modulates the connectivity of the motor engram in specific ways.

As summarized in Figure 8, both the timing with respect to learning and the focality of stimulation influenced the direction of the effects. Transcranial DC stimulation applied before learning showed generally weak modulatory effects. When administered concurrently with learning, the focality of stimulation played a critical role: Targeted anodal tDCS increased the engram’s connectivity, but targeted cathodal or uniform tDCS regardless of the polarity reduced it in an intensity-dependent manner. Given that the engram’s final connectivity is positively correlated with its evoked response [80], such wide-sense “associative” effects of tDCS on motor learning have been observed before. We also predicted with our simulations that, when applied immediately after learning, uniform tDCS generally reduced the connectivity of the engram, but targeted tDCS showed reversed polarity-dependent modulatory effects: anodal tDCS weakens the memory engram, but cathodal stimulation strengthens it. Furthermore, unfocused tDCS affected the connectivity within the memory engram non-uniformly. In summary, our homeostatic plasticity model provides a systematic framework to account for all the confounding factors and mixed results regarding the interaction between tDCS and learning.

**Figure 8:**
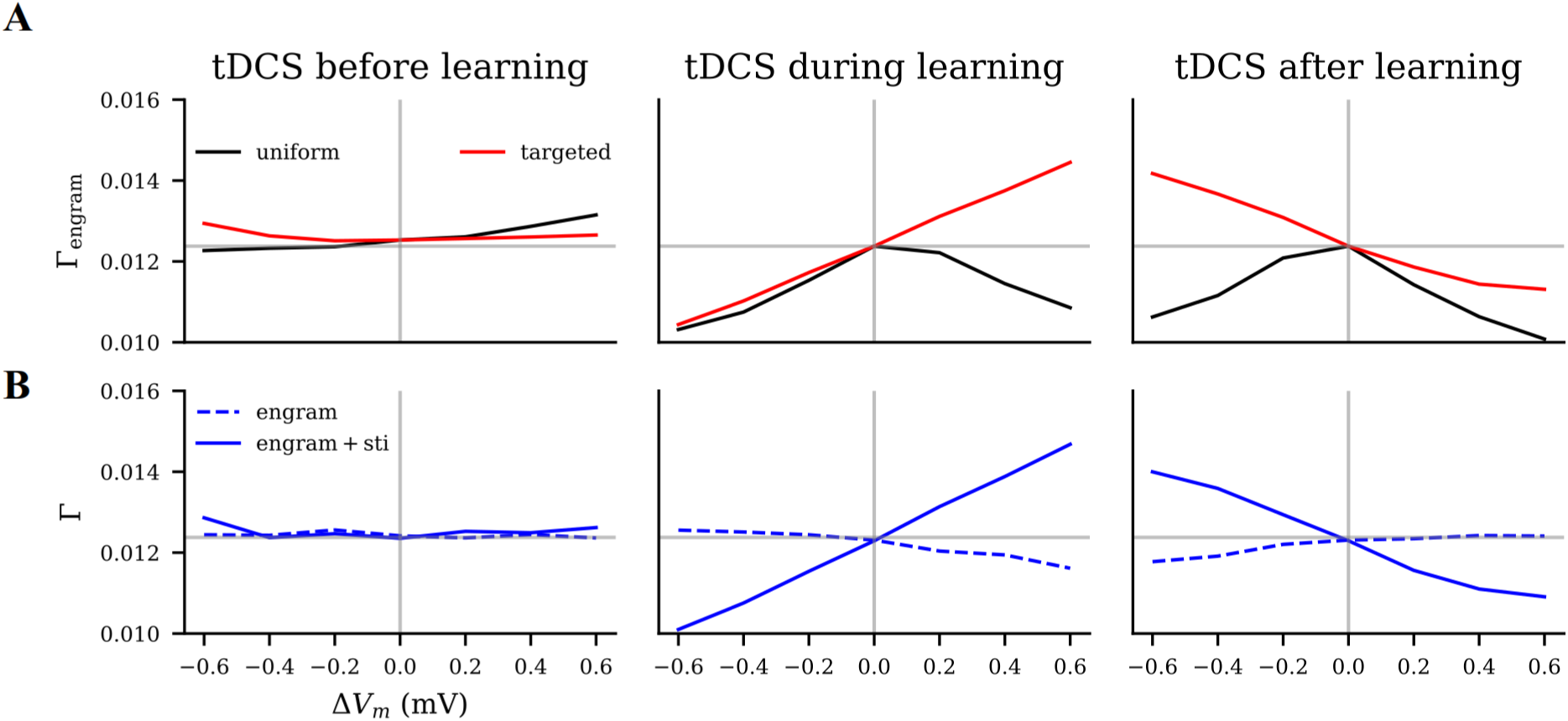
Modulatory effects of tDCS on motor learning as predicted by our model. We estimated the effects of tDCS by the connectivity of the motor engram (Γ_engram_). The x-axis (Δ*V*_m_) denotes the polarity and intensity of the applied DC. The horizontal solid lines (light gray) indicate the engram’s baseline connectivity without tDCS. **A** Effects of targeted and uniform tDCS on engram connectivity when applied immediately before, during, or immediately after learning. **B** Connectivity of the overlapped half and the non-overlapped half of the motor engram, when tDCS with different polarities and intensities was applied either before, during, or after learning.

### Importance of focality and electrode montage

Large electrodes were confirmed to induce more focal electric field distribution than small button-like electrodes used in the high-definition montage [62]. However, the definition of focality that we discussed in the current study is a relevant concept with regard to the neural population engaged by a certain task. We classified it by the degree of overlap that the stimulated neuron population has with the neuron ensemble that encodes a motor program. We were able to show that uniform stimulation systematically reduces the effect of learning, regardless of the relative timing of the application. Targeted stimulation, on the contrary, either boosts or reduces the impact of learning in a predictable way, depending on the current polarity and phase of the stimulus. Our results confirm once again that the focality of the stimulus is critical to achieving a substantial modulatory effect of tDCS [63, 75, 84, 85]. Clearly, the three classes of montage considered in our present work do not cover all variants used in motor learning experiments. Unilateral, bilateral, and high-definition electrode layouts are commonly used [60, 62], and result in mixed effects on motor tasks and learning phases. For example, unilateral or bilateral tDCS applied concurrently increased precision in a follow-up experiment, irrespective of the electrode montage [60]. In contrast, stimulation before learning using a bilateral montage facilitated learning more than unilateral montages [61]. HD-tDCS, which supports focused stimulation, achieved better effects in the clinical treatment of major depressive disorder. Its boosting effects on motor learning do not appear to surpass the conventional unilateral montage [62]. In fact, we were able to confirm that focality generally interferes with the polarity and timing of transcranial stimulation. However, our simple concept disregarded the stimulation of interhemispheric connections, which could be essential to understand the specific properties of a bilateral montage [60].

### Relative intensity of tDCS and motor learning

In human applications, tDCS is widely accepted as a weak and sub-threshold stimulation method [59]. This is the reason we assumed that tDCS provides a weaker stimulus than motor learning in our study. Our model suggests that the relative strength of tDCS compared to motor learning input is critical.

We started with weak transcranial DC stimulation (maximally at 0.6 mV) compared to motor learning (equivalent to 1.5 mV). As suggested for the example of unfocused populations (Figure 5), a strong motor engram is weakened by a weak depolarizing stimulation following it, but not strongly influenced by a weak stimulus applied before it.

However, targeted or uniform tDCS of either polarity applied before learning presented weak but facilitating effects (Figure 8A, left panel), which seems to contradict the diminishing effects of tDCS applied before motor learning observed in human experiments [39, 64, 65, 66, 67, 73]. Consequently, we examine what happens if the transcranial stimulus is stronger than the motor learning input (*±*2.8 mV) in an unfocused montage (Figure 7). Indeed, strong tDCS applied before or after learning distorted the motor engram’s connectivity pattern. Although strong tDCS enhanced the overlapped engram’s connectivity, the ultimate learning performance that may count on the whole motor engram does not necessarily improve. Strongly tDCS which fails to exclusively cover the motor engram will eventually distort the memory information.

### Relative timing of tDCS and motor learning

Another factor confirmed in our model to mediate the modulatory effects of tDCS is the relative timing of tDCS applied to motor learning.

Such results have been frequently observed in human experiments and have been summarized as state-dependent stimulation effects [50, 86, 87]. For example, in the case of pre-learning tDCS, anodal stimulation reduced the learning rate, while cathodal stimulation showed facilitation or impairment [39, 64, 65, 66, 67, 74]. Further studies showed that the impairment caused by pre-learning anodal tDCS was accompanied by a homeostatic increase of GABA_A_ activity [88]. On the basis of these observations, homeostatic plasticity gained popularity as a potential explanation. The idea is that excitability-enhancing tDCS triggers homeostatic inhibition and impairs task performance [74]. On the other hand, concurrent tDCS showed associative effects: Anodal tDCS usually results in improved learning, while cathodal tDCS does not achieve similar results [28, 42, 65]. Concurrent anodal tDCS improves the learning effects not only when applied during the performance of a motor task, but also during observational learning of motor tasks [89]. Such boosting effects of simultaneous anodal tDCS were mostly discussed as non-homeostatic phenomena, as they could not be explained by homeostatic models [90, 91].

Compared to other proposed homeostatic or Hebbian models, our HSP model can consistently explain both types of phenomena. Anodal pre-learning tDCS may hinder learning, when applied uniformly, or at an intensity that out-weights motor input. This is considered a homeostatic phenomenon. In contrast, it is considered a Hebbian phenomenon that concurrent anodal tDCS could facilitate learning. In addition, our model provides a systematic explanation for contradictory or confusing results in tDCS studies with different protocols. The electrode montage of tDCS, for instance, influences the relative coverage of the target region by the electric field, and this different focality can induce opposite effects, even if the polarity and intensity of the stimulus remain the same.

### Memory encoding and consolidation

In addition to pre-learning and concurrent stimulation, our work also predicted the effects of post-learning tDCS on memory consolidation. Uniform stimulation applied post-learning or strong unfocused stimulation hindered memory consolidation, but targeted cathodal tDCS applied to the same brain region where motor learning occurs should boost memory consolidation.

Experimental results on the memory encoding and consolidation effects of post-learning tDCS are lacking. Some studies reported that tDCS applied to M1 facilitates memory encoding [54] or consolidation [92, 93], where the stimulus was administered concurrently with learning, but not after learning. There are also preliminary results showing that the application of anodal tDCS 4 h after learning increased the recall of the learned movement sequences in humans [94]. However, the target area was the premotor cortex and not M1. Our predictions based on the HSP model wait to be addressed by experiments. However, memory consolidation does not necessarily take place in the same region as motor learning [39]. Learning and post-learning tDCS might modulate other influencing factors that are relevant to memory consolidation but beyond the scope of our study, such as neural oscillations during sleep [95].

## Conclusions and outlook

In the current study, we confirmed again that focal tDCS applied in a targeted manner leads to more predictable and reliable modulatory effects. We also showed that the HSP model can explain both the homeostatic and Hebbian plasticity phenomena observed in experiments using tDCS to modulate motor learning. When applied in a targeted manner or with high intensity, pre-learning anodal stimulation could impair the learning process, whereas concurrent anodal stimulation boosted learning. In addition, we predicted that post-learning cathodal tDCS might facilitate learning, which waits to be addressed in future experiments. Compared to previously proposed plasticity models, the HSP model, which is based on homeostatically controlled spine turnover dynamics, provides a more comprehensive explanation and leads to new predictions about the dynamic interaction between tDCS and motor learning.

But can the HSP rule also explain the tDCS effects observed in other forms of learning? Yes and no. The predictions of our model, such as improved learning for targeted concurrent anodal tDCS, match the results found in verbal learning [96] and in working memory tasks [97]. Compared to motor tasks, verbal learning and working memory require the coordination of more brain regions and demand a model to reflect inter-network communication. Our present study is a first attempt to study the effect of transcranial brain stimulation on complex cognitive processes. To bring more insight to this field, more animal studies are still needed to calibrate the cellular mechanism of tDCS and optimize the homeostatic structural plasticity model.

## Supportive Information

### Author contributions

**HL**: conceptualization, methodology, software, validation, formal analysis, investigation, visualization, writing the original draft. **LF**: conceptualization, visualization, writing review & editing. **CN**: conceptualization, visualization, writing review & editing, funding acquisition. **SR**: conceptualization, methodology, writing review & editing, supervision, project administration, funding acquisition.

## Acknowledgments

Additional support by the German Research Foundation (DFG) through EXC 1086, and by the state of Baden-Württemberg through bwHPC and the German Research Foundation (DFG) through INST 39/963-1 FUGG is acknowledged. The authors thank Júlia V. Gallinaro for establishing the fundamental HSP model. We also thank Uwe Grauer from the Bernstein Center Freiburg, as well as Bernd Wiebelt and Michael Janczyk from the Freiburg University Computing Center for their assistance with HPC applications.

