## Supplementary materials for "Resolving inconsistent effects of tDCS on learning using a homeostatic structural plasticity model"

### Neuron model

The membrane potential dynamics of the linear integrate-and-fire (LIF) neuron model is explained by Equation S.1,

$$\tau_m \frac{\mathrm{d}}{\mathrm{dt}} V_i(t) = -V_i(t) + \tau_m \sum_j J_{ij} S_j(t-d) + J_{\mathrm{ML}} S_{\mathrm{ML}}(t-d) + \Delta V(t), \tag{S.1}$$

where $V_i(t)$ is the membrane potential of the neuron $i$, with a resting value at 0 mV. $\tau_m$ is the membrane time constant. $S_j(t) = \sum_k \delta(t - t_j^k)$ represents the spike train generated by the neuron $j$, where $t_j^k$ represents the individual spike times, and $d$ is the synaptic transmission delay. The amplitude of the postsynaptic potential induced in the neuron $i$ upon arrival of a spike from the neuron $j$ is governed by matrix $J_{ij}$. $S_{\mathrm{ML}}(t-d)$ represents the motor learning inputs to a subset of excitatory neurons with synaptic weight $J_{\mathrm{ML}}$. $\Delta V(t)$ represents a membrane polarization imposed by an external electric field. When the membrane potential $V_i(t)$ reaches the threshold $V_{\mathrm{th}}$, an action potential is emitted, and the membrane potential is reset to $V_{\mathrm{reset}} = 10\,\mathrm{mV}$. All parameters of neuron model are summarized in Supplementary Table S1.

### Network model

We modeled the M1 area as an inhibitory-dominated recurrent neural network [1], comprising 10 000 excitatory and 2 500 inhibitory neurons. All synapses involving inhibitory neurons (I-E, inhibitory to excitatory neurons; I-I, inhibitory to inhibitory neurons; E-I, excitatory to inhibitory neurons) were static without plasticity. Each inhibitory neuron randomly received synapses from 10% of the inhibitory and excitatory neurons in the network. Each excitatory neuron also randomly received synapses from 10% of the inhibitory neurons. The static excitatory and inhibitory synapses have a fixed synaptic weight $J_E = 0.1\,\mathrm{mV}$ and $J_I = -0.8\,\mathrm{mV}$. Recurrent synapses among excitatory neurons (E-E) were grown from zero based on the linear HSP rule. Each neuron in the network received Poissonian external input at a rate of $r_{\text{ext}} = 30\,\text{kHz}$. After the 750 s growth period, the network automatically entered an asynchronous-irregular state [1]. All network parameters are listed in Supplementary Table S2.

**Supplementary Table S1:** Parameters of neuron model

| $\tau_m$ | $t_{\text{ref}}$ | $V_0$ | $V_{\text{reset}}$ | $V_{\text{th}}$ |
|---|---|---|---|---|
| 10.0 ms | 2.0 ms | 0.0 mV | 10.0 mV | 20.0 mV |

**Supplementary Table S2:** Parameters for network model

| $N_E$ | $N_I$ | $\Gamma_{E-I}$ | $\Gamma_{I-E}$ | $\Gamma_{I-I}$ | $J_E$ | $J_I$ | $r_{\text{ext}}$ | $J_{\text{ML}}$ | $r_{\text{ML}}$ |
|---|---|---|---|---|---|---|---|---|---|
| 10 000 | 2 500 | 10% | 10% | 10% | 0.1 mV | −0.8 mV | 30 kHz | 0.1 mV | 1.5 kHz |

**Supplementary Table S3:** Parameters for the structural plasticity model

| $\epsilon$ | $\nu$ | $\tau_{\text{Ca}}$ | $\beta_{\text{Ca}}$ |
|---|---|---|---|
| 0.008 | $0.004\,\text{s}^{-1}$ | 10 s | 0.0001 |

### Homeostatic structural plasticity (HSP) rule

The growth or elimination of the synapses of all E-E connections was subject to a rate-based homeostatic structural plasticity rule (HSP) [2]. According to the concept of HSP, E-E connections were formed by presynaptic and postsynaptic elements (boutons and spines). The growth and retraction of the synaptic elements was regulated by the activity-related intracellular calcium concentration ($C(t) = [\text{Ca}^{2+}]$) of individual neurons. Calcium dynamics is summarized in Equation S.2: whenever the neuron emits a spike, the intracellular calcium concentration experiences an influx ($\beta_{\text{Ca}}$) and decays exponentially between spikes with the time constant $\tau_{\text{Ca}}$.

$$\frac{\mathrm{d}}{\mathrm{dt}}C(t) = -\frac{1}{\tau_{\text{Ca}}}C(t) + \beta_{\text{Ca}}S(t) \tag{S.2}$$

Neurons constantly monitored the deviation of their calcium concentration, or in other words, their neural activity, to the equilibrium level (*set-point*). When neural activity falls below the set point, the neuron grows new synaptic elements and forms functional synapses. In contrast, if the firing rate goes above the set point, synapses are broken up, and synaptic elements are removed, leaving the compartment of the broken synapses free for forming synapses again. We adopted a linear growth rule for both the presynaptic and postsynaptic elements:

$$\frac{\mathrm{d}}{\mathrm{dt}} z(t) = \nu \big[1 - \frac{1}{\epsilon} C(t)\big], \tag{S.3}$$

where $z(t)$ is the total number of synaptic elements, $\nu$ is the growth rate, and $\epsilon$ is the set point of calcium concentration. Neurons form functional synapses based on the availability of free synaptic elements, and any pair of neurons can form multiple synapses with the same synaptic weight ($J_E = 0.1\,\mathrm{mV}$). All parameters related to the structural plasticity rule are summarized in the Supplementary Table S3.
